# Evaluation of a One Health mass dog rabies vaccination campaign in a resource-limited urban setting: Evidence from Techiman, Ghana

**DOI:** 10.64898/2026.02.24.707664

**Authors:** Prince Kyere Dwaah, Sylvia Afriyie Squire, Nana Yaa Awua-Boateng, Denis Dekugmen Yar, David Kando, Abraham Num, Helen Djang-Fordjour, Princess Amissah

## Abstract

**Background:** Dog-mediated rabies remains endemic in many parts of sub-Saharan Africa despite the availability of effective vaccines. Achieving and sustaining at least 70% vaccination coverage in dog populations is widely recognized as essential for interrupting transmission; however, there is limited operational evidence from municipal campaigns conducted in rapidly urbanizing, resource-constrained settings. We evaluated the implementation and outcomes of a One Health-coordinated free mass dog rabies vaccination campaign conducted in September 2024 in Techiman South Metropolitan, Ghana.

**Methodology:** A mixed-methods post-campaign assessment was undertaken two weeks after campaign completion. Quantitative data were collected from 200 dog-owning households using structured questionnaires, and administrative vaccination records were reviewed to estimate overall coverage. In addition, key informant interviews were conducted with veterinary officers, public health professionals, environmental health personnel, and community leaders involved in campaign delivery. Descriptive statistics were used to summarize household participation and awareness indicators, while qualitative data were analyzed thematically.

**Principal Findings:** Among surveyed households, 59.5% owned more than one dog and 51.5% reported vaccinating at least one dog during the campaign. Awareness that rabies is fatal increased from 45% (self-reported pre-campaign recall) to 75% after the campaign period, while awareness that rabies is preventable increased from 40% to 70%. Administrative records indicated that approximately 5,600 dogs were vaccinated, corresponding to an estimated 74% metropolitan-level coverage based on projected dog population figures. Stakeholders identified strong multisectoral collaboration and World Rabies Day mobilization as key facilitators. Reported challenges included vaccine supply delays, limited cold-chain capacity, inadequate access for peri-urban communities, and the absence of strategies to vaccinate free-roaming dogs.

**Conclusions/Significance:** These findings suggest that short-duration, one-health mass dog vaccination campaigns can achieve target coverage thresholds in resource-limited urban contexts. However, reliance on projected denominators and exclusion of free-roaming dogs may overestimate effective population immunity. Strengthening monitoring systems, decentralizing access, and incorporating structured strategies for free-roaming dog vaccination will be critical for sustaining rabies control and advancing toward elimination goals.

**AUTHOR SUMMARY:** Rabies is almost always fatal once symptoms begin, yet it can be prevented through vaccination of dogs, which are responsible for nearly all human infections in Africa. Global strategies call for vaccinating at least 70% of dogs each year to stop transmission, but there is limited evidence on how well this works in fast-growing African cities with limited resources.

We evaluated a free mass dog rabies vaccination campaign carried out in Techiman, Ghana, using a One Health approach that brought together veterinary services, public health authorities, and community leaders. We interviewed 200 dog-owning households and reviewed official vaccination records. We also spoke with frontline staff involved in the campaign.

Official records suggested that roughly three-quarters of the estimated dog population was vaccinated, meeting the recommended coverage threshold. Many households reported improved knowledge about rabies following the campaign. However, not all dog-owning households participated, and free-roaming dogs were not systematically vaccinated. Logistical challenges, including vaccine supply delays and limited access in peri-urban communities, were also identified.

Our findings show that coordinated municipal campaigns can reach high vaccination numbers even in resource-limited settings. At the same time, accurate monitoring and inclusion of free-roaming dogs are essential to ensure that reported coverage reflects true protection. Strengthening these components will help Ghana and similar countries move closer to eliminating dog-mediated human rabies.

## INTRODUCTION

Rabies is an acute viral zoonosis that remains endemic in many parts of Africa and Asia, causing an estimated 59,000 human deaths annually, with more than 95% attributable to dog-mediated transmission (Hampson et al., 2015; WHO, 2023a). Despite the availability of safe and effective vaccines for both humans and animals, rabies continues to impose a substantial public health burden in low- and middle-income countries (LMICs), where access to post-exposure prophylaxis (PEP), surveillance capacity, and sustained animal vaccination programs are often limited (Abela-Ridder et al., 2016; WHO, 2023b). Children and underserved populations are disproportionately affected, reflecting inequities in health care access, dog ownership practices, and animal health service coverage (Hampson et al., 2015).

From an epidemiological perspective, sustained interruption of dog-mediated rabies transmission is achievable through high annual vaccination coverage in domestic dog populations. Both mathematical modelling and empirical field studies indicate that vaccinating at least 70% of the dog population can interrupt transmission and reduce spillover into human populations, even in settings with high dog density and substantial free-roaming dog populations (Lembo et al., 2010; WHO et al., 2018). Consequently, mass dog vaccination has been identified as the most cost-effective rabies prevention strategy in multiple endemic settings, compared with reliance on human PEP alone (Zinsstag et al., 2009; Del Rio Vilas et al., 2022).

Despite this evidence base, implementing mass dog vaccination at scale in LMICs remains challenging. Common barriers include incomplete dog population estimates, limited veterinary workforce capacity, cold-chain and vaccine supply constraints, heterogeneous community participation, and difficulties accessing peri-urban and free-roaming dog populations (Abela-Ridder et al., 2016; Mbilo et al., 2021). Reported vaccination coverage therefore often relies on administrative data that do not fully account for unregistered or free-roaming dogs and may consequently overestimate true population-level protection. These challenges highlight the importance of independent post-campaign evaluations that combine household-level surveys with qualitative assessments of program implementation (Mbilo et al., 2021; WHO, 2023c).

In response to persistent global rabies transmission, the World Health Organization, together with the Food and Agriculture Organization, the World Organisation for Animal Health, and UNICEF, launched the “Zero by 30” strategic framework to eliminate human deaths from dog-mediated rabies by 2030 (WHO et al., 2018). Central to this framework is a One Health approach that integrates veterinary services, public health systems, and community engagement to strengthen vaccination delivery, surveillance, and awareness. While national-level progress has been documented in selected regions, evidence from subnational and municipal vaccination campaigns particularly in rapidly urbanizing, resource-limited settings remains limited (Del Rio Vilas et al., 2022).

Ghana continues to experience endemic rabies transmission, with recurrent reports of dog bites and sporadic human rabies cases across multiple regions (Turkson and Wi-Afedzi, 2020; Tasiame et al., 2021). Dog vaccination coverage remains variable and is frequently constrained by financial barriers, limited enforcement of dog ownership regulations, and low public awareness (Punguyire et al., 2017; Turkson and Wi-Afedzi, 2020). Techiman South Municipality, a rapidly growing commercial and transport hub with high human–dog contact rates in the Bono East Region, has been identified as a recurrent hotspot for dog bites and confirmed human rabies cases, reflecting the interaction of dense human populations, free-roaming dogs, and constrained veterinary infrastructure (Punguyire et al., 2017; Azanduna & Quandzie, 2024).

To address these challenges, a One Health–coordinated free mass dog rabies vaccination campaign was implemented in Techiman South Municipality in September 2024, aligned with World Rabies Day activities. The campaign sought to improve access to vaccination services, increase public awareness, and achieve vaccination coverage consistent with World Health Organization recommendations. Understanding the operational performance of such campaigns is essential for informing future rabies control strategies, particularly in urban and peri-urban LMIC settings where program sustainability and equitable coverage remain critical challenges.

The objective of this study was to evaluate the vaccination coverage, community reach, and implementation challenges of the Techiman mass dog rabies vaccination campaign using a mixed-methods post-campaign assessment. By integrating household survey data, administrative vaccination records, and qualitative insights from key stakeholders, this study provides operational and implementation-level evidence to inform the design, monitoring, and evaluation of mass dog vaccination campaigns in comparable urban LMIC settings.

## MATERIALS AND METHODS

### Study area

The study was conducted in Techiman South Metropolitan, located in the Bono East Region of Ghana. Techiman is a rapidly urbanizing commercial and transport hub with an estimated population of approximately 173,000 residents (Ghana Statistical Service, 2021). The metropolitan comprises urban, peri-urban, and adjoining rural communities and supports substantial free-roaming and owned dog populations. Recurrent reports of dog bites and confirmed human rabies cases have been documented in the municipality, indicating a sustained risk of rabies transmission (Punguyire et al., 2017; Azanduna & Quandzie, 2024).

### Description of the One Health Rabies Vaccination Campaign

The mass dog rabies vaccination campaign was implemented in September 2024 in Techiman South Metropolis using a One Health framework that integrated veterinary, public health, and community-level actions. The campaign was jointly coordinated by the Veterinary Services Directorate and the Metropolitan Health Directorate, with active involvement of environmental health officers, community leaders, and local information services. Veterinary teams were responsible for vaccine delivery, cold-chain management, dog handling, and vaccination, while public health personnel supported risk communication, community sensitization, and linkage of dog-bite victims to appropriate health facilities for post-exposure prophylaxis. Community leaders and local media facilitated mobilization, dissemination of campaign messages, and identification of vaccination sites. Campaign activities were aligned with World Rabies Day, leveraging global messaging to enhance public trust and participation. This multisectoral collaboration aimed to simultaneously reduce rabies risk at the animal source, improve community awareness of rabies prevention, and strengthen coordination between animal and human health services, consistent with internationally recommended One Health approaches to rabies control.

### Study design

A mixed-methods, cross-sectional post-campaign evaluation was conducted in November 2024, approximately two weeks after completion of a free mass dog rabies vaccination campaign. Although the evaluation was implemented after the campaign, limited pre-campaign information was collected retrospectively to provide contextual baseline data.

Specifically, the household questionnaire included recall-based items assessing respondents’ knowledge and awareness before the campaign, including awareness that rabies is fatal, awareness that rabies is preventable, and whether respondents had previously heard of rabies vaccination activities. These items were framed explicitly as “before the campaign” questions and were used to generate descriptive pre-campaign indicators.

Post-campaign data captured household participation in vaccination, exposure to campaign communication activities, and current rabies knowledge. Pre- and post-campaign indicators were compared descriptively to assess changes in awareness associated with the campaign period. No causal inference was intended, and findings are interpreted as indicative of perceived changes rather than measured baseline-to-endline effects.

### Study population and sampling

The quantitative component targeted dog-owning households residing in communities included in the vaccination campaign. Eligible participants were adults (≥18 years) who resided in the municipality during the campaign period and reported ownership of at least one dog.

Communities were selected purposively based on campaign implementation coverage, followed by snowball sampling to identify dog-owning households. This approach was selected to assess campaign reach and operational performance rather than to estimate population-level prevalence. One eligible adult per household was interviewed.

### Sample size

The target sample size was calculated using Cochran’s formula for cross-sectional studies (Cochran, 1977), assuming a vaccination proportion of 50%, a 95% confidence level, and a 7% margin of error, yielding a minimum required sample of 196 households. To account for incomplete responses, a final sample of 200 dog-owning households was included.

### Data collection

#### Quantitative data

A structured, pretested questionnaire was administered through face-to-face interviews. The instrument captured household demographics, number of dogs owned, vaccination status during the campaign, rabies knowledge, and exposure to campaign communication activities. The questionnaire was piloted in neighbouring municipalities, namely Techiman North, Wenchi, and Sunyani West, with ten respondents in each municipality, before deployment and was revised for clarity.

#### >Qualitative data

Key informant interviews (KIIs) were conducted with ten stakeholders, including veterinary officers, public health professionals, and community leaders involved in campaign planning or implementation. Interviews explored coordination mechanisms, community response, logistical constraints, and gaps in vaccination coverage. Interviews were conducted using a semi-structured guide and audio-recorded with participant consent.

#### Administrative data

Administrative vaccination records were obtained from the Techiman Veterinary Services Directorate, including data from the Metropolitan Veterinary Clinic and the Disease Investigation Farm/Regional Veterinary Laboratory. These records were used to estimate the total number of dogs vaccinated during the campaign.

Estimated vaccination coverage was calculated using projected dog population size derived from municipal veterinary reports and the World Health Organization–recommended human-to-dog ratio of 5:1. These estimates are presented as indicative measures of coverage rather than precise population-level estimates.

#### Data analysis

Quantitative data were entered into Microsoft Excel and analyzed using SPSS version 26. Descriptive statistics, including frequencies and percentages, were calculated to summarize household vaccination participation and rabies awareness indicators.

Qualitative data were analyzed using thematic analysis following the six-phase framework described by Braun and Clarke 2006. Interview transcripts were reviewed, coded inductively, and analyzed to identify recurrent themes related to campaign facilitators and implementation barriers. Findings from qualitative and quantitative components were triangulated during interpretation.

#### Ethical considerations

The study involved no medical intervention and collected no personal identifiers. Formal institutional ethical approval was not obtained. Administrative permission was granted by the Regional Veterinary Office (Bono East Region), the Metropolitan Veterinary Services Directorate and the Metropolitan Health Directorate. Written informed consent was obtained from all participants before data collection.

## RESULTS

### Household dog ownership and vaccination participation

A total of 200 dog-owning households participated in the survey. Among these, 81 households (40.5%) reported owning one dog, while 119 households (59.5%) owned more than one dog, as shown in *Table 1*. From *Figure 1*, overall, 103 households (51.5%) reported vaccinating at least one dog during the vaccination campaign.

**Table 1:** Household dog ownership among surveyed households.

| Number of dogs owned | Frequency | Percentage |
| --- | --- | --- |
| 1 | 81 | 40.5 |
| More than 1 | 119 | 59.5 |

**Footnote:** Percentages represent the proportion of surveyed households reporting each ownership category.

**Table 2.** Rabies knowledge and awareness before and after the campaign.

| Variable | Pre-Campaign (%) | Post-Campaign (%) |
| --- | --- | --- |
| Awareness that rabies is fatal | 45 | 75 |
| Awareness that rabies is preventable | 40 | 70 |
| Heard of the rabies vaccination campaign | 38 | 95 |
| First information source: Radio/TV | 35 | 38 |
| First information source: Health workers | 28 | 32 |
| First information source: Peer social networks (community meetings, friends, family) | 37 | 45 |

**Footnote:** Values represent the percentage of respondents affirming each indicator at each survey time point.

**Figure 1:**
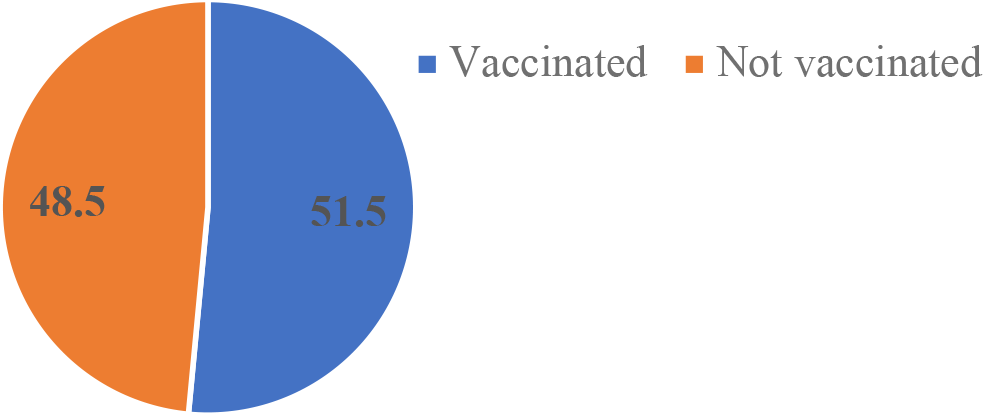
Proportion of dog-owning households reporting vaccination of at least one dog during the 2024 mass rabies vaccination campaign

Among households that participated in vaccination, the number of dogs vaccinated per household varied according to dog ownership size. Households owning multiple dogs were more likely to report partial vaccination of their dogs than complete household-level coverage.

### Community awareness of rabies

Before the campaign, 90 respondents (45.0%) reported awareness that rabies is fatal, and 80 respondents (40.0%) reported awareness that rabies is preventable. Following the campaign period, 150 respondents (75.0%) reported awareness that rabies is fatal, and 140 respondents (70.0%) reported awareness that rabies is preventable. Also, 95% of the respondents reported having heard of the vaccination campaign. The most commonly reported initial sources of information were radio or television (35.0%), health workers (28.0%), and peer or community networks, including family members and community meetings (37.0%). Following the campaign, reported reliance on radio or television and health workers increased modestly, while reliance on peer networks declined.

### Administrative vaccination data and estimated coverage

Administrative records from the Techiman Veterinary Services Directorate indicated that approximately 5,600/8,000 dogs were vaccinated during the campaign across participating communities in the metropolitan. Based on projected dog population estimates derived from municipal veterinary reports and application of the World Health Organization– recommended human-to-dog ratio of 5:1, this corresponds to an estimated vaccination coverage of approximately 74% at the metropolitan level.

Administrative records did not include stratification by ownership status or vaccination of free-roaming dogs, and coverage estimates were therefore limited to owned dogs presented at vaccination points.

### Key informant interview findings

Ten key informant interviews were completed with three veterinary officers, two public health officials, two environmental health officers and three community leaders. Four recurrent themes emerged from the thematic analysis, as shown in *Table 3*.

**Table 3.** Key Informant Themes from WRD Campaign Evaluation.

| Theme | Description | Illustrative Quote |
| --- | --- | --- |
| Enhanced Community Engagement Through WRD | WRD created momentum, community trust, and higher participation. | People came out in large numbers because they heard it was a global event. They respected the initiative more. (Veterinary Officers, Environmental Health Officers and Public Health Officers) |
| Operational Challenges During Campaign Implementation | Vaccine supply delays, limited cold-chain capacity, and long queues discouraged some participants from attending. | There were delays in vaccine supply, and the queues were very long. (Veterinary Officer) |
| Inaccessibility for Rural and Peri-Urban Communities | A lack of transportation and distant sites limited access for outlying areas. | Many people from outlying communities wanted to bring their dogs, but there was no transportation or nearby site. (Community leaders) |
| Gaps in Stray Dog Coverage and Control | Absence of Catch-Vaccinate-Release (CVR) teams prevented coverage of stray dogs. | We could not vaccinate stray dogs; without CVR, herd immunity is threatened. (Veterinary Officers, Environmental Health Officers and Public Health Officers) |

## DISCUSSION

This study provides an implementation-focused evaluation of a One Health mass dog rabies vaccination campaign conducted in a resource-limited urban setting in Ghana. The findings demonstrate that substantial numbers of dogs can be vaccinated during short-duration campaigns; however, important gaps remain in achieving uniform and epidemiologically robust coverage.

Administrative data indicated an estimated vaccination coverage of approximately 75%, calculated using projected dog population estimates. Although this estimate exceeds the threshold commonly cited for interruption of rabies transmission, the exclusion of free-roaming dogs and reliance on population projections suggest that true population-level immunity may be lower (Lembo et al., 2010; Mbilo et al., 2021; WHO, 2023a). Studies in similar African contexts have shown that failure to vaccinate free-roaming dogs can sustain rabies transmission despite high reported coverage among owned dogs (Hampson et al., 2015; Mbilo et al., 2021).

The discrepancy between administrative coverage estimates and household-reported participation observed in this study is consistent with findings from other LMIC settings, where administrative records tend to overestimate effective coverage due to incomplete denominators and aggregation of vaccination counts (Mbilo et al., 2021; WHO, 2023b). Household-level surveys provide complementary insights into accessibility and utilization of services but may underestimate coverage when households vaccinate dogs outside sampled communities or when partial household vaccination occurs (Abela-Ridder et al., 2016). Together, these observations underscore the importance of triangulating administrative and household data when assessing vaccination program performance.

Community awareness of rabies was moderate before the campaign and higher following the campaign period. Although causal attribution cannot be inferred from this cross-sectional evaluation, similar increases in rabies knowledge following mass vaccination activities and awareness campaigns have been reported elsewhere (Lembo et al., 2010; Turkson and Wi-Afedzi, 2020). The prominence of mass media and health workers as information sources aligns with previous studies in Ghana and other African settings but may insufficiently reach peri-urban or marginalized populations without complementary community-based engagement strategies (Tasiame et al., 2021; WHO, 2023c).

Qualitative findings highlighted World Rabies Day activities as an important mechanism for community mobilization, consistent with evidence that globally recognized health events can enhance participation by increasing visibility and trust (Del Rio Vilas et al., 2022). Operational challenges identified in this study, including logistical constraints, limited cold-chain capacity, and restricted geographic access, have been widely reported as barriers to campaign-based rabies control in LMICs (Abela-Ridder et al., 2016; Mbilo et al., 2021). These constraints are particularly relevant in peri-urban settings, where population density and mobility complicate centralized service delivery.

The absence of strategies to vaccinate free-roaming dogs was a critical limitation of the campaign. Catch-vaccinate-release (CVR) approaches and other dog population management strategies have been shown to improve coverage and sustain transmission interruption when integrated into mass vaccination programs (Lembo et al., 2010; WHO et al., 2018). Without systematic inclusion of free-roaming dogs, vaccination campaigns may achieve high administrative coverage without fully addressing the epidemiological reservoir (Hampson et al., 2015; Mbilo et al., 2021).

This evaluation also illustrates the operational value of a One Health approach at the municipal level. Coordination between veterinary services, public health authorities, and community leadership supported campaign delivery and information dissemination. Similar multisectoral approaches have been identified as critical for effective rabies control, particularly where veterinary and public health systems face resource constraints (Del Rio Vilas et al., 2022; WHO et al., 2018). However, sustainability will depend on institutionalized coordination, predictable financing, and workforce capacity beyond episodic campaign activities.

Several limitations should be considered when interpreting these findings. The cross-sectional design precludes causal inference regarding changes in awareness or behaviour. Household data were self-reported and may be subject to recall bias. Administrative vaccination data lacked ownership status and geospatial disaggregation, limiting precision in coverage estimation. Finally, free-roaming dogs were not directly assessed, constraining the evaluation of true population-level immunity.

Despite these limitations, this study contributes operational evidence from a subnational urban setting, an area where empirical data remain limited. The findings support the role of mass dog vaccination as a central component of rabies control while highlighting the need for improved monitoring systems, decentralized access and systematic inclusion of free-roaming dogs to sustain progress toward rabies prevention.

## CONCLUSION

This study demonstrates that mass dog vaccination campaigns implemented through multisectoral collaboration can achieve substantial vaccination outputs in resource-limited urban settings. Administrative records indicated high campaign reach, while household-level data highlighted important gaps in access and participation, particularly among multi-dog households and peri-urban communities. These findings underscore the need to interpret vaccination coverage estimates cautiously and to triangulate administrative and household data when evaluating program performance.

The absence of structured strategies to vaccinate free-roaming dogs represents a critical limitation for sustained rabies control. Without systematic inclusion of these populations, high administrative coverage among owned dogs may not translate into sufficient population-level immunity. Logistical constraints, including vaccine supply timing, cold-chain limitations, and geographic accessibility, further influenced campaign implementation and should be addressed in future planning.

The observed coordination between veterinary services, public health authorities, and community leadership supports the operational feasibility of a One Health approach at the municipal level. Strengthening this collaboration through routine planning, decentralized delivery models, and improved monitoring systems may enhance campaign effectiveness and sustainability.

Overall, these findings highlight that while short-duration mass dog vaccination campaigns can play a central role in rabies prevention, their epidemiological impact depends on accurate population estimates, inclusive vaccination strategies, and sustained multisectoral engagement. Addressing these factors will be essential for advancing rabies control efforts in rapidly urbanizing settings in Ghana and similar contexts.

## AUTHOR’S CONTRIBUTION

All authors contributed equally.

## ACKNOWLEDGMENTS

We acknowledge the support of the Veterinary Services Directorate, Ghana Health Service, Techiman Municipal Assembly, and the dedicated community volunteers. Special thanks to the staff of the Disease Investigation Farm and Regional Veterinary Laboratory, Techiman, for their leadership and commitment throughout the project.

## CONFLICTS OF INTEREST

The authors declare no conflict of interest. The views expressed in this communication are those of the authors and do not necessarily reflect the official policies of the Ghana Health Service, the Ministry of Food and Agriculture or any other affiliated institution.

## FUNDING

This study did not receive any specific grant from funding agencies in the public, commercial or not-for-profit sectors.

## INSTITUTIONAL REVIEW BOARD STATEMENT

This study was conducted in accordance with international research guidelines as outlined by the WHO Ethics Review Committee. Although formal ethical approval was not obtained but permission was sought from the Regional Veterinary Officer of Bono East, the Metropolitan Environmental Health Directorate and the Metropolitan Health Directorate.

## INFORMED CONSENT STATEMENT

Informed verbal and written consent were obtained from all participating dog owners, community members, and healthcare personnel involved in the intervention. Participation was entirely voluntary, and no identifiers were collected. All participants were informed of the purpose, process, benefits, and potential risks of the intervention.

## DATA AVAILABILITY STATEMENT

The data supporting the findings of this study are available from the corresponding author upon reasonable request. Aggregated anonymized data on vaccination coverage is stored at the Disease Investigation farm and Regional Veterinary Laboratory, the Metropolitan Veterinary Office.

## Notes

### Competing Interest Statement

The authors have declared no competing interest.

